# Registration of sugar beet genetic stocks FC308 (PI701378) and FC309 (PI700990)

**DOI:** 10.1101/2023.05.17.541007

**Authors:** Olivia E Todd, Amy L Nielson, Ann Fenwick, Linda E Hanson, Kelley L Richardson, Kevin M Dorn

## Abstract

Sugar beet (*Beta vulgaris* L.) genetic stock lines FC308 (PI701378) and FC309 (PI700990) are two highly homozygous and phenotypically homogenous sources of resistance to two sugar beet pests. FC308 is resistant to sugar beet cyst nematode (*Heterodera schachtii)*, but susceptible to Fusarium yellows, while FC309 is Fusarium yellows resistant, but sugar beet cyst nematode susceptible. These two populations were developed by the USDA-ARS in Fort Collins, CO, derived from unreleased USDA-ARS Salinas germplasm lines. Multiple field and greenhouse trials for Fusarium yellows and nematode resistance confirmed the uniform phenotypes of both lines for each disease. To enable molecular breeders to rapidly screen proprietary markers in silico, gene space assemblies for each line were also developed. Genome sequencing of individual plants from each line, as well as pooled sequencing of sub-populations of both lines indicated fixation of a previously reported genetic variant linked to sugar beet cyst nematode resistance in FC308. However, two previously reported variants linked to Fusarium resistance appear to be unlinked to the resistance found in FC309. Collectively, these two new genetic stocks should prove useful as sources of resistance to Fusarium yellows and sugar beet cyst nematode, as model genetic backgrounds for studying plant-pathogen interactions, and for mapping the resistance genes present in these lines.

## 1. INTRODUCTION

American beet sugar production accounts for 55-60% of the sugar produced domestically, representing a $1+ billion/year industry (USDA-ERS, 2021). A multitude of biotic and abiotic stressors continually threaten both yield and post-harvest losses in sugar beet, which challenge 7M acres in the United States alone (BSDF, 2021). Members of the *Fusarium oxysporum* species complex (herein called “*Fusarium oxysporum”*) cause a suite of diseases on a variety of host crops, including Fusarium wilt, seedling disease Fusarium stalk blight and Fusarium yellows in sugar beet (Adaba, 1994; Hanson & Jacobsen, 2009; Harveson, 2009; Ruppel 1991; Stewart, 1931). *Fusarium* remains a critical challenge to beet sugar production in the United States, impacting 79,900 acres, causing an estimated $9.6M USD in lost value to the American beet sugar market in 2021 (BSDF, 2021). *Fusarium oxysporum* has been reported to have cross-pathogenicity and is able to infect multiple host plant species (Hanson, Hill, Jacobsen, & Panella, 2009; Martyn, Rush, Biles, & Baker, 1989). These diseases often cause total loss of the infected sugar beet plant and result in necrosis of both above and below ground biomass. Resistant germplasm is in high demand by regional growers and is one of the best methods to combat disease caused by *F. oxysporum* (Fravel, Olivain, & Alabouvette, 2003). Information for marker assisted selection of *F. oxysporum* resistance has been developed in cereal crops such as wheat (Anderson, 2007) and specialty crops including tomato, chick pea, melon and cabbage (Ahmad, Mumtaz, Ghafoor, Ali, & Nisar, 2014; El Mohtar et al., 2007; Gao, Liu, Zhu, & Luan, 2015; Liu et al., 2017). Marker-trait associations in sugar beet breeding are notoriously lacking in the public sector across most target disease resistance traits. Only one recent study developed surrogate molecular markers to enable marker assisted selection for *Fusarium* resistance in sugar beet (De Lucchi et al., 2017). Across beet crops, there are no functionally validated resistance genes against *Fusarium*-induced diseases.

Sugar beet cyst nematode (*Heterodera schachtii)* (SBCN) is increasing in prevalence throughout the world and is a cause for concern for sugar beet growers. SBCN infection can have many effects on the crop, including yield loss, stunting, chlorosis and wilting (Cooke, 1991; Lilley, Atkinson, & Urwin, 2005) Many molecular studies have been conducted to elucidate plant resistance mechanisms to SBCN in *Beta vulgaris* including the application of ascorbate oxidase, an enzyme involved in oxidation of apoplastic ascorbic acid leading to induced systemic resistance in sugar beet (Singh, Nobleza, Demeestere, & Kyndt, 2020). The Singh et al. study, as well as others, highlight the importance of phytohormones in SBCN resistance and defense (Ghaemi et al., 2020). Other research has explored the efficacy of resistant transgenes from both sweet potato in the hairy root system and other crop wild relatives such as *Patellifolia procumbens* (Cai et al., 2003; Kumar et al., 2021). Several SBCN resistant lines with unknown resistance mechanisms have been released in affiliation with the USDA-ARS (Lewellen, 1995; Richardson, 2012, 2018, 2022).

## 2. METHODS

### 2.1 Population Development

The USDA-ARS Sugar Beet Genetics Lab in Fort Collins, Colorado has been screening wild germplasm with the goal of identifying and developing pathogen resistant germplasm for nearly 60 years. USDA-ARS sugar beet breeder Dr. Bob Lewellen developed sister lines Salinas 7927-4-309 for its resistance to Fusarium stalk blight and susceptibility to sugar beet cyst nematode and Salinas 7927-4-308 for resistance to sugar beet cyst nematode and susceptibility to Fusarium yellows. Later, mapping populations were made with these two lines (Richardson, 2022). Both Salinas 7927-4-309 and Salinas 7927-4-308 were developed from C927-4 (PI 628756), originally variable response to sugar beet cyst nematode (Richardson, 2022). Seed of Salinas 7927-4-309 and Salinas 7927-4-308 was received from the Salinas, CA ARS group in 2011.

A greenhouse-based bulk increase of 44 total inter-pollinated roots from the 7927-4-308 seed lot created the 20131014 seed lot, which was subsequently evaluated for resistance to SBCN (Table 2). An additional bulk increase of the 20131014 seed lot, consisting of 261 total inter-pollinated plants gave rise to the 20151015 seed lot and herein referred to as FC308 (PI701378). An alternative, genetically bottlenecked population (20202523) derived from Salinas 7927-4-308 was also developed using a greenhouse-based inter-pollination of 25 individual plants. Deep short-read sequencing of pooled DNA from tissue from each of these 25 plants, which were a part of another large-scale population re-sequencing project (NCBI BioProject PRJNA563463, SRR12806811), was also utilized in this study.

The Salinas 7927-4-309 seed lot was bulk increased in a greenhouse environment, producing the 20131013 seed lot which was later evaluated for SBCN resistance in both field and greenhouse tests, and found to be susceptible (Table 2). Four plants from the Salinas 7927-4-309 seed lot were identified as possessing strong resistance to *Fusarium oxysporum* in a greenhouse screen. These plants were vernalized, and subsequently allowed to self-pollinate under high intensity LEDs in HEPA-filtered growth chambers for single seed descent. The progeny seed of the individual plant that produced the most seed, 20211021-4s, was planted in a field isolation plot at the Colorado State University Agricultural Research, Development, and Education Center (Fort Collins, Colorado USA) in 2021. At the end of the season, 97 mother roots were harvested, vernalized for 120 days, then planted in the greenhouse in the spring of 2022 to make the 20221005 seed lot, herein referred to as FC309 (PI700990) (Figure 1A). Similar to FC308, an alternative population bottlenecked through an inter-pollination of 25 plants from the Salinas 7927-4-309 seed lot (20202524) were subjected to pooled DNA sequencing (NCBI Bioproject PRJNA563463, SRR12806809).

**Figure 1.**
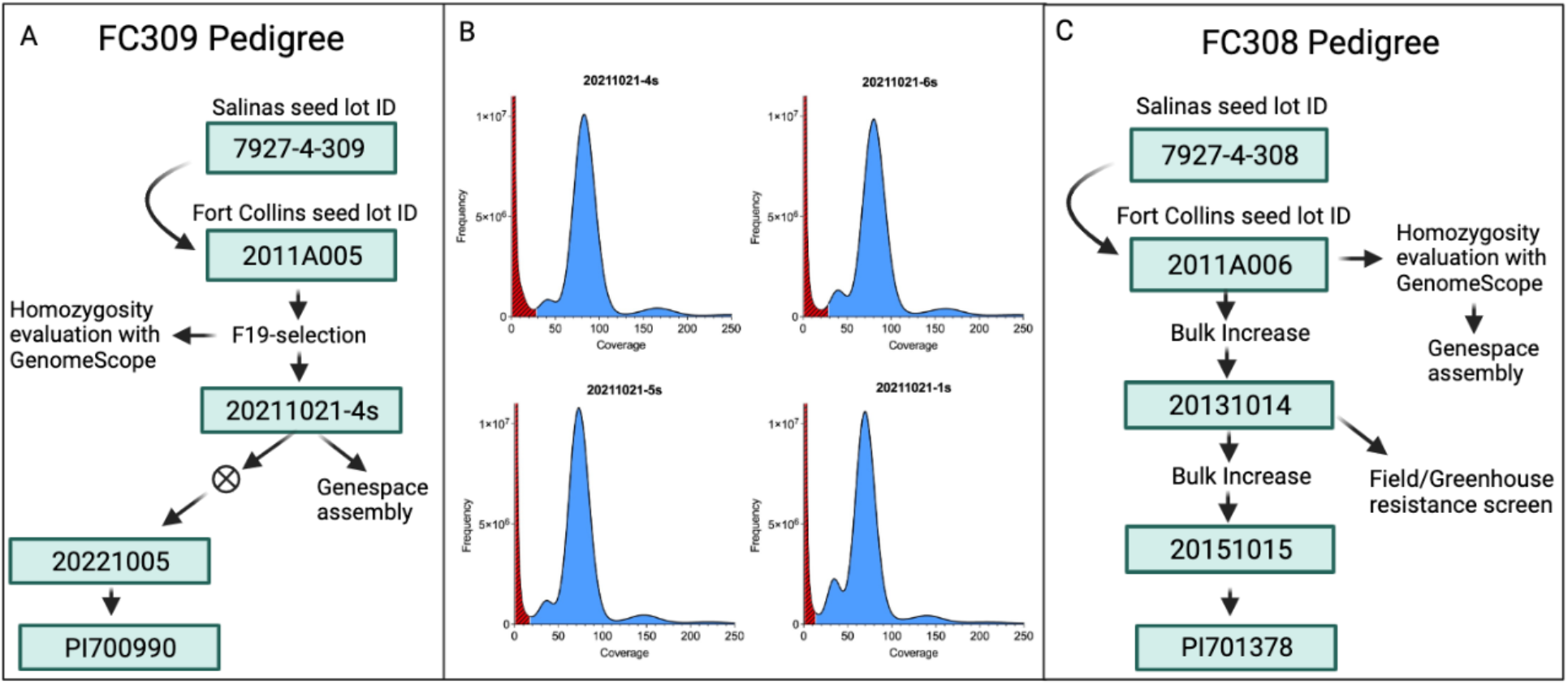
A. PI700990 line generation beginning with Salinas 7927-4-309. Upon receipt in Fort Collins, CO, it was given the 2011A005 designation and screened with *Fusarium oxysporum* strain F19. Six surviving plants were selected for self-pollination and sequencing, but only 20211021-4s was selected for seed increase and line release. B. Histogram of k-mer counts showing genome properties of *Fusarium* resistance selected lines 20211021-4s (top left), 20211021-6s (top right), 20211021-5s (bottom left) and 20211021-1s (bottom right). Using a k-mer length of 21 determined by the software Jellyfish, the GenomeScope models determine read error rate from the Illumina sequencing platform (highlighted in red crosshatch and in the “Error” column of Table 1), and an estimate of unique, nonrepetitive sequencing data and average sequence duplication rate (highlighted in solid blue and shown in Table 1). C. PI701378 line generation beginning with requesting 7927-4-308 seed from Salinas, CA. Upon receipt in Fort Collins, CO, it was given the 2011A006 designation and bulk increased in the greenhouse in 2013. An additional bulk increase was made in the field in 2015. Field tests for sugar beet cyst nematode resistance were conducted with the 20131014 seed, and shown to be not statistically different than the resistant cultivar check (Table 2).

### 2.2 Greenhouse Fusarium Screening

All of the *Fusarium* greenhouse screening was conducted with an 18h photoperiod, a mean daytime temperature of 26C and a mean night time temperature of 18C. The relative humidity was 30-40%. Plants from line FC309 were seeded into six-inch pots. When plants were four to six weeks old, beets were uprooted from their pots, washed in a tub of clean water and submerged in a suspension of *Fusarium oxysporum* strain F-19 (10^6^ CFU/ml) for five minutes, and then repotted. Treated plants were evaluated based on a 0-4 disease severity scale, where 0 indicates a healthy plant 21-30 days after inoculation (Hanson & Hill, 2004). Surviving plants were vernalized and then self-pollinated.

**Table 1:** Various genome structural characteristics for four *Fusarium oxysporum* strain F19 selected sugar beet lines following model iteration over the histogram file generated by the software program Jellyfish. Notably, heterozygosity is low in all individuals from 20211021 (FC309).

| Line | Heterozygosity | Unique Sequences | Error | Duplication |
| --- | --- | --- | --- | --- |
| 20211021-1s | 0.33% | 70.4% | 0.21% | 0.76% |
| 20211021-4s* | 0.14% | 71.8% | 0.24% | 0.79% |
| 20211021-5s | 0.18% | 71.0% | 0.20% | 0.74% |
| 20211021-6s | 0.27% | 71.5% | 0.19% | 0.87% |

\*Gene space assembly produced using Illumina reads (USDA\_Bvulg\_FC309\_v0.1), described in methods.

**Table 2:** Least squares mean of sugarbeet cyst nematode (SBCN) canopy appearance score over four years at the Imperial Valley Research Center in Brawly, CA. Lines 7927-4-308 and 7927-4-309 are progenitor lines to FC309 and FC309, respectively. Beta 8520N is SBCN resistant, C37 is SBCN susceptible.

| Line | SBCN canopy score (1-5)* |  |  |  |
| --- | --- | --- | --- | --- |
|  | 2009 | 2011 | 2012 | 2013 |
| 7927-4-308 | 2.49 de | 2.99 cde | 3.23 cd | 2.30 def |
| 7927-4-309 | 4.16 ab | 4.89 a | 3.12 cd | 3.82 abc |
| Beta 8520N | 2.11 ef | 1.45 f | 2.22 ef | 3.15 bcde |
| C37 | 3.88 abc | 3.87 abc | 2.96 cde | 1.62 f |

\*SBCN canopy score 1 = little or no chlorosis and necrosis of the leaves and approximately 90 to 100% of leaves are living; 3 = approximately 50% of leaves are living and 50% show either necrosis or are dead; 5 = most or all leaves are dead or nearly dead; within and between years, lines with the same letter are not significantly different at $p>0.05$ .

### 2.3 Field Screening for Resistance to Sugar beet cyst nematode

Trials were planted at the Imperial Valley Research Center in Brawley, CA under conditions favoring SBCN (Lewellen and Pakish 2005). The field contained 4.2 ha of SBCN-infested soil. Each year, tests were rotated throughout this space to ensure no direct replanting for 2 years. Ten years of rotated sugar beet plantings increased the nematode populations to high levels (Becker et al., 1996). Tests were conducted in 2009, 2011, 2012, and 2013. Seed of FC308 and FC309 progenitors 7927-4-308, 7927-4-309 respectively, Beta 8520N (SBCN resistant), and C37 (PI 590715, SBCN susceptible) were planted in September in a completely randomized design of one-row, 4-m-long plots with eight replications. To assess SBCN resistance under field conditions, the canopies were scored 9 months after planting and rated from 1 to 5 on the basis of appearance (Lewellen and Pakish 2005). A score of 1 indicates little or no chlorosis or necrosis of the leaves; approximately 90 to 100% of leaves are living and are similar to the same material in companion trials without infestation to SBCN. A score of 3 means approximately 50% of leaves are living and 50% are either necrotic or dead. A score of 5 indicates most or all of the leaves are dead or nearly dead or that there is evidence that there had been leaves but these are now deteriorated. Under the combined effects of high nematode populations and very high daytime temperatures (average 42°C), SBCN-infested plants reach a critical wilting point from which they do not recover (Lewellen and Pakish 2005). Canopy appearance scores are an indicator of relative SBCN resistance. Analysis of variance of canopy appearance scores was performed with the SAS MIXED procedure (ver. 9.3, SAS Institute, Inc.) with line as a fixed effect and the random statement with years as random effects. Variance components were estimated using the REML method and Satterthwaite degrees of freedom. Mean separation was calculated using least squares means (LSMeans statement in SAS). A multiple comparison test, the Tukey-Kramer method, was performed (p < 0.05). There was a significant line × year interaction (*p* < 0.0001). LSMean canopy appearance scores are presented for each year (Table 2). In 2009, 2011, and 2013, LSMean SBCN canopy appearance scores for 7927-4-308 were significantly lower than 7927-4-309; in 2012, they were not significantly different. In 2009 and 2013, 7927-4-308 scores were not significantly different than the resistant check, Beta 8520N, but they were significantly higher in 2011 and 2012.

The FC308 (20131014) and FC309 (20131013) seed lots were screened for resistance to sugar beet cyst nematode in Johnstown, CO in a field plot with endemic SBCN pressure. The nursery consisted of 20 entries, including a commercial hybrid resistant and susceptible check with 4 replications arranged in a completely randomized design. Two-row, 16 feet long plots on 22 inch beds were used at a seeding rate of 2.5 grams of seed/single row plot. The field was planted May 1^st^, 2015, and rated August 5^th^ and 28^th^, 2015. A visual rating scale of 1-5 was used to rate severity. Pairwise comparisons with a Kramer-Tukey adjustment were used to compare entries to the resistant and susceptible checks (Table 3).

**Table 3:** Sugar beet cyst nematode evaluation for number of cysts per sugar beet line per location between 5 years. Resistant and susceptible checks varied across sites. The FC308 line 20131014 is derived from the Fort Collins seed lot 2011A006 (Figure 1c). The FC309 line 20131013 is derived from the Fort Collins seed lot 2011A005.

| Line | Field | Greenhouse Sites |  |
| --- | --- | --- | --- |
|  | Johnstown,<br>Colorado 2015 | 2020 | 2021 |
| Resistant check | 1 | 0.5 | 4 |
| 20131014 (308) | 2<sup>a</sup> | 7.69 | 17.829 |
| 20131013 (309) | 3.5 | 34.4 | 45.37 |
| Susceptible check | 3.25 |  |  |

<sup>a</sup>Line not significantly different than the resistance check based on a pairwise comparison with Kramer-Tukey adjustment ( $\alpha= 0.05$ ).

### 2.4 Greenhouse Screening for Resistance to Sugar beet cyst nematode

The 20131013 (FC309) and 20131014 (FC308) seed lots were screened in a greenhouse-based SBCN resistance test. Greenhouse screening consisted of planting seeds into 12 cm diameter pots filled with washed sand. Single seedlings were transplanted into tubes (15 cm x 20 cm x 120 cm) filled with a soil-sand mixture 10 days after planting. Tubes were placed in plastic boxes (30 cm x 20 cm x 13 cm) for the remainder of the experiment. A total of 20 seedlings per line (5 plants x 4 replicates) were inoculated with ca. 500 sugar beet cyst nematodes suspended in 3 mL of water. Plants were grown in a mean temperature of 25C with a 16h photoperiod, and 70% relative humidity. Four weeks following inoculation, plants were removed from tubes and washed with tap water. The number of individual cysts per plant were manually counted. The student’s t-test was used (90% confidence interval) compare the average number of cysts per repetition (data not shown).

### 2.5 Illumina Sequencing for Homozygosity Estimation with GenomeScope

Four individual plants from the 2011A005 seed lot (FC309) that exhibited resistance in the *F. oxysporum* F19 greenhouse screening and one 2011A006 (FC308) individual were selected for whole genome sequencing with the Illumina platform (Figure 1A and C). DNA was isolated from each plant using the Qiagen DNeasy Plant Mini Kit. Illumina paired end sequencing resulted in an average of 67 gigabases (Gb) of reads per FC309 individual, or approximately 90X genome coverage and 22.1 Gb (29.5X coverage) was generated for the FC308 individual. Raw sequencing reads were trimmed and filtered for quality and adaptor contamination using BBduk, a part of the BBMap package (v38) using the flags “k=23 hdist=1 ktrim=r qtrim=r trimq=20 mink=11 tpe tbo” using the ‘adapters.fa’ file included in the BBMap release for adapter filtering/trimming. Homozygosity of each of the four FC309 and one FC308 was evaluated using GenomeScope (Vurture et al., 2017). Using the count program, we first evaluated k-mer frequency with Jellyfish2 (v2.2.9) with the flags -C -m 21 -s 2000000000. The k-mer distribution was exported using the jellyfish histo function (Figure 1B). All of the sequenced plants were grown after vernalization under high intensity LEDs in HEPA-filtered growth chambers for single seed descent.

### 2.6 Validation of Previously Published Molecular Markers

Gene-space assemblies, which are assemblies that capture a majority of the gene-encoding portion of the genome, were created using the trimmed and filtered Illumina reads from one FC309 individual (20211021_4s) and one FC308 individual (2011A006_7s). Reads were assembled using a de-Bruijn graph strategy in the De Novo Assembly Toolkit in CLC Genomics Workbench (v20.0.4) using the following parameters: Automatic Word Size, Automatic Bubble Size, Minimum Contig Length:500, Auto-detect paired distances, Perform scaffolding, Map reads back to contigs (mismatch cost: 2, insertion cost: 3, deletion cost: 3, length fraction:0.5, similarity fraction:0.8). Contaminating sequences identified by the NCBI Foreign Contamination Screen tool (https://github.com/ncbi/fcs) were removed from both of the assemblies. The final gene space assemblies were named “USDA_Bvulg_FC309_v0.1” and “USDA_Bvulg_FC308_v0.1”. Details for access are under “Data Availability”.

To identify the corresponding genome position of the molecular marker linked to the sugar beet cyst nematode resistance HsBvm-1 (Stevanato et al., 2015), the amplicon from SNP192 (5’-AATCATATATCAACACCTATAACGCATCTAAATGACAGGGTATACAACTACCAGGCCACATATATAGCCAAACAGCTCAAGCCTGTACAAAAGACTAAACA-3’) was searched against both the FC308 (USDA_Bvulg_FC308_v0.1) and FC309 (USDA_Bvulg_FC309_v0.1) gene space assemblies using BLASTn (v2.9.0) implemented in CLC Genomics Workbench (BLASTn parameters: Match=2, Mismatch=3, Existence=5, Extension=2, Expectation Value = 10, Word size = 11, Mask lower case = No, Mask low complexity regions = Yes). Corresponding positions for this marker sequence were identified and extracted from both the FC308 (USDA_Bvulg_FC308_v0.1) and FC309 (USDA_Bvulg_FC309_v0.1) assemblies. The HsBvm-1 SNP192 marker sequence was aligned with this region from both assemblies using CLC Genomics Workbench (v20.0.4, Create Alignment 1.02, Gap open cost=10, Gap extension cost = 1, End gap cost = As any other, Alignment mode = Very accurate (slow), Redo alignments=No, Use fixpoints=No).

To evaluate the allelic status of previously published molecular markers linked to *Fusarium* resistance (De Lucchi et al., 2017) and sugar beet cyst nematode (Stevanato et al., 2015) in sugar beet, Illumina DNA sequencing reads derived from pools of 25 individual plants for both FC308 (BioProject PRJNA563463, SRR12806811) and FC309 (BioProject PRJNA563463, SRR12806809) were utilized, along with the USDA_Bvulg_FC308_v0.1 and USDA_Bvulg_FC309_v0.1 gene space assemblies. To find the homologs of the *Fusarium* resistance gene analogs identified by De Lucchi et al. (2017) in each gene space assembly, the predicted protein sequences for the RefBeet v1.1 gene models Bv2_043450_zhxk and Bv7_171470_ojty (obtained from https://bvseq.boku.ac.at/Genome/Download/RefBeet-1.1/ in file <u>RefBeet.genes.1302.pep.fa</u>) were searched against USDA_Bvulg_FC308_v0.1 and USDA_Bvulg_FC309_v0.1 gene space assemblies using tBLASTn (Expectation value = 10.0, Word size = 3, Mask lower case = No, Mask low complexity regions = Yes, Protein Matrix and Gap Costs = BLOSUM62_Existence_11_Extension_1). The homologous contigs for the target RGA markers were identified in both gene space assemblies (Bv2_043450: USDA_Bvulg_FC308_v0.1_contig_0000294 and USDA_Bvulg_FC309_v0.1_contig_0000649,

Bv7_171470: USDA_Bvulg_FC309_v0.1_contig_0018888, USDA_Bvulg_FC308_v0.1_contig_0027168).

The pooled sequencing reads for FC309 (SRR12806809) and FC308 (SRR12806811) were mapped to both gene space assemblies using ‘Map Reads to Reference’ tool (v1.7) in CLC Genomics Workbench v 20.0.4 (Masking mode=No masking, Match score = 1, Mismatch cost = 2, Cost of insertions and deletions – Linear gap cost, Insertion cost=3, Deletion cost = 3, Length fraction = 0.9, Similarity Fraction = 0.9, Global alignment = No, Auto-detect paired distances = Yes, Non-specific match handling = Ignore).

## 3. CHARACTERISTICS

### 3.1 Whole genome homozygosity

High depth Illumina sequencing of the FC309 progenitor population identified a single plant (20211021-4s) with the highest k-mer derived global homozygosity inferred from GenomeScope (Figure 1B), which was selected for single seed decent. Similarly, moderate depth Illumina sequencing of a single plant from the FC308 progenitor population indicated high genome-wide homozygosity.

### 3.2 Fixation at known molecular markers

Corresponding positions for the HsBvm-1 SNP192 marker were identified in both the v0.1 gene space assemblies for FC308 (USDA_Bvulg_FC308_v0.1_contig_0000996 (position 20,344 bp – 20,444 bp), e-value of 7.03e-44, bit score = 178.92) and FC309 (USDA_Bvulg_FC309_v0.1_contig_0000796, position 15,245 bp – 15,345 bp, e-value of 7.89 e-44, bit score = 178.92) (data not shown). The corresponding FC308 contig contained the ‘G’ allele for SNP192 marker position, whereas the FC309 contig contained the ‘C’ allele. This result is consistent with the results from De Lucchi et al. (2017), with the exception of an additional variant 42 base pairs downstream of the SNP192 variant position (T/A, data not shown). Manual examination of the mapping of the pooled sequencing of 25 individual FC308 plants mapped against the USDA_Bvulg_FC308_v0.1 reference revealed 98.38% read support (61/62 reads) for the ‘G’ allele (SBCN resistant) at the SNP192 position. For FC309, there was 100% support (60/60 reads) for the ‘C’ allele (SBCN susceptible) at the SNP192 position.

The gene space assembly tBLASTn best hits for both the Bv2_043450 and Bv7_171470 Fusarium markers from De Lucchi et al. clearly identified the corresponding sequences from both FC308 and FC309. Interestingly, the there was no sequence variation at the Bv2_043450 marker in the gene space assembly contigs. The pooled sequencing datasets from each line mapped back to their respective gene space assemblies indicated that the ‘A’ (Fusarium resistant) allele was nearly fixed in both lines, with 38/38 read support at position 17,365 of USDA_Bvulg_FC308_v0.1_contig_0000294 and 51/52 read support at position 11,044 of USDA_Bvulg_FC309_v0.1_contig_0000649. The results were similar for the Bv7_171470 SNP marker. For FC309, 60/63 reads supported the ‘A’ (resistant) allele at position 2,937 of USDA_Bvulg_FC309_v0.1_contig_0018888, whereas 62/63 reads supporting the ‘A’ (resistant) allele at position 744 of USDA_Bvulg_FC308_v0.1_contig_0027168). Collectively, these results point to the Fusarium resistance found in FC309 is not linked to the markers described in De Lucchi et al.

### 3.3 Fusarium Resistance

FC309 exhibits strong, uniform resistance to *Fusarium oxysporum* strain F-19, as demonstrated in multiple greenhouse inoculation trials (data not shown). Upon treatment with the root dip protocol previously described, there are no yellow or necrotic leaf developments throughout a 30-day period post inoculation, and disease severity is zero (Figure 2). Conversely, FC308 has uniform susceptibility to *Fusarium oxysporum* strain F-19. FC308 has the potential to serve as a susceptible check in Fusarium nurseries, greenhouse screens, and as a model genotype for studying the host-pathogen interaction.

**Figure 2.**
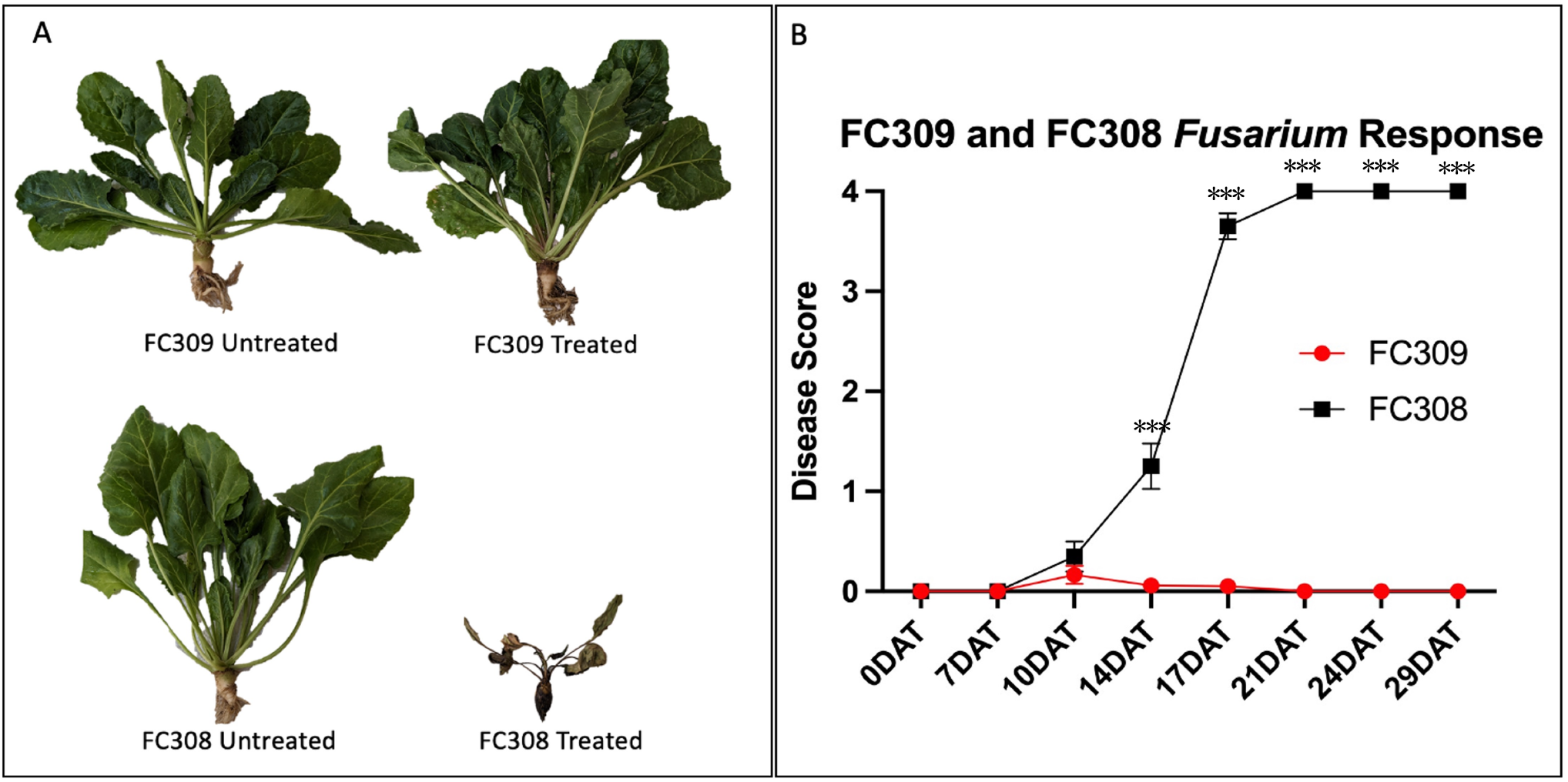
A. PI700990 (FC309) (top) and PI701378 (FC308) (bottom) phenotype response to *Fusarium oxysporum* F19 0 and 30 days after treatment (DAT). Roots are significantly deteriorated and white fungal hyphae can be seen on FC308 as the plant necrotizes what it left of the tissue. B. Time course of FC309 and FC308 disease response to *Fusarium oxysporum* strain F19 rated according to the 0-4 Hanson et al. (2004) scale. *Asterisks represent statistical significance according to an unpaired t-test at each time point between FC309 and FC308 at p = 0.0001.

### 3.4 Sugar beet cyst nematode Resistance

FC308 exhibits uniform tolerance to *Heterodera schachtii* as evidenced by field and greenhouse trials (Table 2). The FC308 seed lot 20131014 exhibited statistically similar SBCN scores to the commercial resistant check in the 2015 Johnstown, CO nursery, whereas the FC309 seed lot 20131013 exhibited susceptibility similar to the susceptible check line. In two separate greenhouse screens, FC308 exhibited SBCN tolerance (Table 2), but with worse SBCN ratings (number of cysts) compared to the CN65-9 CMS line (seed lot 2007A097) which contains the Hs1^Pro1^ translocation from *Patellifolia procumbens* (Kumar et al, 2021).

### 3.5 General Phenotypic Characteristics

Both FC308 and FC309 are self-fertile, however, FC309 is exceptionally fecund in the greenhouse, with single plant self-pollination producing up to 50 grams per plant. Both FC308 and FC309 are segregating for hypocotyl color (21:6 and 33:20 red to green, respectively). The status of monogerm for FC308 and FC309 are 23% and 54% respectively. FC308 germinates at approximately 17 sprouts per gram of seed, and FC309 germinates at approximately 73 sprouts per gram of seed.

## 4. AVAILABILITY

### 4.1 PI700990 and PI701378 Genetic Stocks

Both genetic stocks are currently available as PI700990 (FC309) and PI701378 (FC308) and are maintained at the National Lab for Genetic Resource Preservation (NLGRP) in Fort Collins, CO. Larger quantities for both seed lots will be maintained by the USDA-ARS Sugar Beet Genetics Lab for distribution upon written request to the corresponding author. These genetic stocks can be searched for and ordered from NPGS using GRIN-Global.

## 5. DATA AVAILABILITY STATEMENT

Genome sequencing datasets described in this study have been deposited at the National Center for Biotechnology Information (NCBI) Sequence Read Archive (SRA) under BioProject PRJNA880324 for USDA_Bvulg_FC309_v0.1 and BioProject PRJNA923373 for USDA_Bvulg_FC308_v0.1. Illumina genome assemblies for both lines are available at NCBI under <Pending GenBank Accession Number> for USDA_Bvulg_FC309_v0.1 and < Pending GenBank Accession Number > for USDA_Bvulg_FC308_v0.1. The pooled sequencing data from 25 individual plants for both lines are available under NCBI BioProject PRJNA563463 (FC309: SRR12806809, FC308: SRR12806811).

## ABBREVIATIONS

FY: Fusarium yellows
CFU: colony-forming units
DAI: days after inoculation
F. SP.: Forma specialis
NLGRP: National Lab for Genetic Resource Preservation
SBCN: Sugar beet cyst nematode
USDA-ARS: United States Department of Agriculture – Agricultural Research Service

## AUTHOR CONTRIBUTIONS

OET: Investigation, writing original draft, review and editing; ALN: Methodology, writing original draft, review and editing; AF: Investigation and editing; KMD: Conceptualization, investigation, writing original draft, review and editing, funding acquisition, supervision; LH: Resources and Conceptualization; KR: Resources, Investigation, and Conceptualization.

## ACKNOWLEDGMENTS

The authors acknowledge Dr. Bob Lewellen and Dr. Lee Panella, and all the members of their respective USDA-ARS labs, for their contribution of to the development of this germplasm. We also acknowledge the Beet Sugar Development Foundation and the Western Sugar Cooperative Joint Research Committee for contributing funding for this project.

## CONFLICT OF INTEREST

The authors declare no conflict of interest.

## Notes

### Competing Interest Statement

The authors have declared no competing interest.

## REFERENCES

Abada, K.A. (1994), Fungi causing damping-off and root rot on sugar beet and their biological control with Trichoderma harzianum. Agriculture Ecosystems and Environment. 51:333=337

Ahmad, Z., Mumtaz, A. S., Ghafoor, A., Ali, A., & Nisar, M. (2014). Marker assisted selection (MAS) for chickpea Fusarium oxysporum wilt resistant genotypes using PCR based molecular markers. Mol Biol Rep, 41(10), 6755–6762. doi:10.1007/s11033-014-3561-3

Anderson, J. A. (2007). Marker-assisted selection for Fusarium head blight resistance in wheat. Int J Food Microbiol, 119(1-2), 51–53. doi:10.1016/j.ijfoodmicro.2007.07.025

Becker, J.O., Wrona, A.F., and Lewellen, R.T. (1996) Effect of solarization and soil fumigation on sugarbeet cyst nematode population, 1993–95. Biol. Cult. Tests Control Plant Dis. 11:19.

BSDF, Beet Sugar Development Foundation (2021). Production Challenges Survey

Cai, D. G., Thurau, T., Tian, Y. Y., Lange, T., Yeh, K. W., & Jung, C. (2003). Sporaminmediated resistance to beet cyst nematodes (Heterodera schachtii Schm.) is dependent on trypsin inhibitory activity in sugar beet (Beta vulgaris L.) hairy roots. Plant Mol Biol, 51(6), 839–849. doi: 10.1023/A:1023089017906

Cooke, D. A. (1991). The effect of beet cyst nematode, Heterodera schachtii, on the yield of sugar-beet in organic soils. Ann Appl Biol, 118(1), 153-160. doi: 10.1111/j.1744-7348.1991.tb06093.x

De Lucchi, C., Stevanato, P., Hanson, L., McGrath, M., Panella, L., De Biaggi, M., … Concheri, G. (2017). Molecular markers for improving control of soil-borne pathogen Fusarium oxysporum in sugar beet. Euphytica, 213(3), 71

El Mohtar, C. A., Atamian, H. S., Dagher, R. B., Abou-Jawdah, Y., Salus, M. S., & Maxwell, D. P. (2007). Marker-assisted selection of tomato genotypes with the I-2 gene for resistance to Fusarium oxysporum f. sp. lycopersici Race 2. Plant Dis, 91(6), 758–762. doi:10.1094/PDIS-91-6-0758

Fravel, D., Olivain, C., & Alabouvette, C. (2003). Fusarium oxysporum and its biocontrol. New phytol, 157(3), 493–502

Gao, P., Liu, S., Zhu, Q. L., & Luan, F. S. (2015). Marker-assisted selection of Fusarium wilt-resistant and gynoecious melon (Cucumis melo L.). Genet Mol Res, 14(4), 16255–16264. doi:10.4238/2015.December.8.16

Ghaemi, R., Pourjam, E., Safaie, N., Verstraeten, B., Mahmoudi, S. B., Mehrabi, R., … Kyndt, T. (2020). Molecular insights into the compatible and incompatible interactions between sugar beet and the beet cyst nematode. BMC Plant Biol, 20(1). doi: 10.1186/s12870-020-02706-8

Hanson, L., & Hill, A. (2004). Fusarium species causing Fusarium yellows of sugarbeet.

Hanson, L., Hill, A., Jacobsen, B., & Panella, L. (2009). Response of sugarbeet lines to isolates of Fusarium oxysporum f. sp. betae from the United States

Hanson, L.E. (2009). Other Fusarium-associated problems. pp. 121-122 In Compendium of Beet Diseases and Pests (2^nd^ ed.) Harveson, R.M., Hanson, L.E.and Hein, G.L. eds. APS Press, St. Paul, MN

Hanson, L.E. and Jacobsen, B.J. (2009). Fusarium yellows. pp.28–20 In Compendium of Beet Diseases and Pests (2^nd^ ed.) Harveson, R.M., Hanson, L.E.and Hein, G.L. eds. APS Press, St. Paul, MN

Harveson, R.M. (2009). Fusarium root rot. pp. 30–31 In Compendium of Beet Diseases and Pests (2^nd^ ed.). Harveson, R.M., Hanson, L.E.and Hein, G.L. eds. APS Press, St. Paul, MN

Kumar, A., Harloff, H. J., Melzer, S., Leineweber, J., Defant, B., & Jung, C. (2021). A rhomboid-like protease gene from an interspecies translocation confers resistance to cyst nematodes. New phytol, 231(2), 801–813

Lewellen, R. T. (1995). Registration of 3 cyst-nematode resistant sugar-beet germplasms - C603, C603-1, and C604. Crop Sci, 35(4), 1229–1230. doi: 10.2135/cropsci1995.0011183X003500040090x

Lewellen, R.T., and Pakish, L.M. (2005) Performance of sugarbeet cyst nematode resistant cultivars and a search for sources of resistance. In: Proceedings of the American Society of Sugar Beet Technologists, Palm Springs, CA. 25 Mar. 2005. p. 122–123.

Lilley, C. J., Atkinson, H. J., & Urwin, P. E. (2005). Molecular aspects of cyst nematodes. Mol Plant Pathol, 6(6), 577–588

Liu, X., Han, F., Kong, C., Fang, Z., Yang, L., Zhang, Y., … Lv, H. (2017). Rapid introgression of the Fusarium wilt resistance gene into an elite cabbage line through the combined application of a microspore culture, genome background analysis, and disease resistance-specific marker assisted foreground selection. Front Plant Sci, 8, 354. doi:10.3389/fpls.2017.00354

Martyn, R., Rush, C., Biles, C., & Baker, E. (1989). Etiology of a root rot disease of sugar beet in Texas. Plant Dis, 73(11), 879–884

Richardson, K. L. (2012). Registration of sugarbeet germplasm lines CN12-446, CN12-770, and CN72-652 with resistance to sugarbeet cyst nematode. J Plant Regist, 6(2), 216–220

Richardson, K. L. (2018). Registration of sugar beet germplasm lines CN921-515 and CN921-516 with sugar beet cyst nematode resistance from Beta vulgaris subsp maritima. J Plant Regist, 12(2), 264–269. doi:10.3198/jpr2017.09.0062crg

Richardson, K. L. (2022). Registration of sugar beet mapping populations CN239, CN240, and CN241 segregating for resistance to Heterodera schachtii from sea beet. J Plant Regist, 16(2), 459–464

Ruppel, E.G. (1991). Pathogencity of Fusarium spp. from diseased sugar beets and variation among sugar beet isolates of F. oxysporum. Plant Disease. 75:486–489

Singh, R. R., Nobleza, N., Demeestere, K., & Kyndt, T. (2020). Ascorbate oxidase induces systemic resistance in sugar beet against cyst nematode Heterodera schachtii. Frontiers in plant science, 11. doi: 10.3389/fpls.2020.591715

Stevanato, P., Trebbi, D., Panella, L., Richardson, K., Broccanello, C., Pakish, L., … Saccomani, M. (2015). Identification and validation of a SNP marker linked to the gene HsBvm-1 for nematode resistance in sugar beet. Plant Molecular Biology Reporter, 33(3), 474–479

Stewart, D. (1931). Sugar-beet yellows caused by Fusarium conglutinans var. betae. Phytopathology, 21(1), 59–70

USDA-ERS. (2021). U.S. Sugar Production. Retrieved from https://www.ers.usda.gov/topics/crops/sugar-sweeteners/background/

Vurture, G. W., Sedlazeck, F. J., Nattestad, M., Underwood, C. J., Fang, H., Gurtowski, J., & Schatz, M. C. (2017). GenomeScope: fast reference-free genome profiling from short reads. Bioinformatics, 33(14), 2202–2204

